# Considerations on activity determinants of fungal polyphenol oxidases based on mutational and structural studies

**DOI:** 10.1101/2021.02.26.433144

**Authors:** Efstratios Nikolaivits, Alexandros Valmas, Grigorios Dedes, Evangelos Topakas, Maria Dimarogona

**Affiliations:** Industrial Biotechnology & Biocatalysis Group, Biotechnology Laboratory, School of Chemical Engineering, National Technical University of Athens, Athens, Greece; Department of Biology, University of Patras, Patras, Greece; Laboratory of Structural Biology and Biotechnology, Department of Chemical Engineering, University of Patras, Patras, Greece

**Keywords:** polyphenol oxidase, X-ray structure, site-directed mutagenesis, tyrosinase, kinetic analysis, *Thermothelomyces thermophila*, protein engineering

## Abstract

Polyphenol oxidases (PPOs) are an industrially relevant family of enzymes, being involved in the post-harvest browning of fruits and vegetables, as well as in human melanogenesis. Their involvement lies in their ability to oxidize phenolic or polyphenolic compounds, that subsequently form pigments. PPO family includes tyrosinases and catechol oxidases, which in spite of their high structural similarity, exhibit different catalytic activities. Long-standing research efforts have not yet managed to decipher the structural determinants responsible for this differentiation, as every new theory is disproved by a more recent study. In the present work, we combined biochemical along with structural data, in order to rationalize the function of a previously characterized PPO from *Thermothelomyces thermophila* (*Tt*PPO). The crystal structure of a *Tt*PPO variant, determined at 1.55 Å resolution, represents the second known structure of an ascomycete PPO. Kinetic data of structure-guided mutants prove the implication of “gate” residue L306, residue H_B1_+1 (G292) and H_B2_+1 (Y296) in *Tt*PPO function against various substrates. Our findings demonstrate the role of L306 in the accommodation of bulky substrates and that residue H_B1_+1 is unlikely to determine monophenolase activity as suggested from previous studies.

**IMPORTANCE:** PPOs are enzymes of biotechnological interest. They have been extensively studied both biochemically and structurally, with a special focus on the plant-derived counterparts. Even so, explicit description of the molecular determinants of their substrate specificity is still pending. Especially for ascomycete PPOs, only one crystal structure has been determined so far, thus limiting our knowledge on this tree branch of the family. In the present study, we report the second crystal structure of an ascomycete PPO. Combined with site-directed mutagenesis and biochemical studies, we depict the amino acids in the vicinity of the active site that affect enzyme activity, and perform a detailed analysis on a variety of substrates. Our findings improve current understanding of structure-function relations of microbial PPOs, which is a prerequisite for the engineering of biocatalysts of desired properties.

## **1.** Introduction

Polyphenol oxidases (PPOs) are a family of enzymes comprising tyrosinases-TYR (EC 1.14.18.1), catechol oxidases-CO (EC 1.10.3.1) and aureusidin synthases-AUS (EC 1.21.3.6). Genes expressing them can be found in all domains of life from humans, animals and insects to plants, fungi, bacteria and archaea. Their role in nature is basically the synthesis of pigments for protective or other purposes depending on the organism. This takes place by the oxidation of phenolic compounds into quinones, which then undergo non-enzymatic reactions towards the formation of pigments (1, 2). Organisms from different kingdoms use different precursors for their pigments, hence they express enzymes with affinity to these specific compounds. For example, animals synthesize their pigments from nitrogen-bearing phenolic compounds like tyrosine, dopamine and catecholamines, while in plants and fungi the most common melanin precursors are catechol, 1,8-dihydroxynaphthalene and some phenolic acids (eg gallic, caffeic and protocatechuic). Bacteria on the other hand seem to be able to utilize both types of precursors to form their melanins (3).

In this manner, PPOs are classified depending on their substrate specificity to those that hydroxylate tyrosine or other similar monophenolic compounds (TYR-monophenolase activity) and to those who can oxidize catecholic derivatives (TYR/CO/AUS-diphenolase activity). This classification seems to be somewhat outdated, though, since recent studies have shown that previously classified COs possess monophenolase activity for their natural substrates, like in the case of AUS from *Coreopsis grandiflora* (*Cg*AUS) (4). Similarly, the PPO from *Aspergillus oryzae* was initially characterized as CO (*Ao*CO4), because it cannot oxidize tyrosine. However, it seems to be able to hydroxylate small monophenols, as well as diphenols in the *ortho*-position prior to their further oxidation to quinones (5). These recent findings are indicative of the need to reconsider the classification and nomenclature of this enzyme family.

PPOs contain one type-III copper site, which reversibly binds dioxygen and consists of two copper ions (CuA and CuB), each coordinated by three histidine residues (H_A1-3_ and H_B1_-3). Type-III copper site can be found in at least four states: *oxy*-, *met*-, *deoxy*- and *hydroperoxide*-forms. In the *oxy*-form, the two Cu^II^ ions bind a dioxygen molecule, while in the *met*-form, which is considered the resting state, the two tetragonal coppers are bridged by a hydroxide or water molecule (6). Reaction of an *o*-diphenol with the *met*-form results in the reduced *deoxy*-form (Cu^I^-dicopper), which can quickly turn into the active *oxy*-form by binding dioxygen. *Hydroperoxide*- form can be formed either by oxygen binding to the reduced site or by peroxide activation for substrate hydroxylation (7, 8). Monophenols, on the other hand, require the *oxy*-PPO form to start the catalytic cycle (6).

Even though the active-site of all type-III proteins shows high similarity, their substrate scope differs. In the past decade, a lot of effort has been made towards the elucidation of the reaction mechanism of PPOs and the molecular determinants of their ability to hydroxylate monophenols. However, these determinants still remain unclear and every theory is overthrown by a newly discovered enzyme or crystal structure. One of the first theories for the lack of monophenolase activity was the presence of a bulky residue (aka “gate” residue) atop CuA (most commonly Phe in plant COs), which was thought to sterically hinder monophenols from binding to CuA, where the hydroxylation step was thought to take place (9). This theory was disproved when the first crystal structure of a plant TYR was elucidated having the bulky Phe at this position (10). Additionally, the work of Goldfeder et al showed that both substrate types bind between the two metal ions, slightly oriented towards CuA (11), while for *Ao*CO4, which possesses hydroxylase activity, it was shown that upon substrate binding a conformational change of the “gate” residue takes place (5). Another key residue important for the binding of common tyrosinase substrates (like tyramine) was thought to be the one adjacent to the second CuB coordinating histidine (H_B2_+1), with Arg favoring substrate binding (12).

It can be seen that deprotonation of a monophenol is considered to be the crucial initial step in order to begin the monophenolase catalytic cycle, and the base implicated in this step has long been under investigation. The possession of hydroxylase activity has been related to the presence of a water molecule that is coordinated and activated by an Asn at position H_B1_+1 and a highly conserved Glu positioned several residues before H_B1_+1 (11). This water molecule is said to deprotonate the incoming monophenol, which is the first step towards its hydroxylation (13). In the case of a PPO from *Vitis vinifera* (*Vv*PPO), the mutation of Gly at position H_B1_+1 to Asn resulted in increased enzyme activity on tyramine and tyrosol (14). Examples of enzymes that do not have an Asn at this position but can still hydroxylate monophenols (not necessarily tyrosine and tyramine) are *Cg*AUS (Thr), two *Agaricus bisporus* PPOs (Asp), two apple TYRs (Ala and Gly) and *Ao*CO4 (Gly) (8). In fact, a mechanism for the hydroxylation by *Ao*CO4 has been proposed, where the enzyme’s copper site is in *hydroperoxide*-form and monooxygenation occurs by electrophilic aromatic substitution without the need for deprotonation (5). Recently, docking studies revealed that the preference towards diphenolic substrates, exhibited by PPOs bearing a Gly in position H_B1_+1, is explained by a “laying down” orientation of the substrate, which is stereochemically prohibited in PPOs bearing a bulkier Asn residue (15).

In addition to the above, many recent studies point out the His residues that coordinate the active site copper ions as potential proton accepting bases. Matoba et al were the first to unveil that monophenol deprotonation takes place via the active site bound peroxide. They also highlighted the mobility of both copper ions and the release of H_A2_ and H_A3_ from CuA during each catalytic cycle (16, 17). Crystallographic work on *A. oryzae* TYR also pointed out the migration of both copper ions and the detachment of H_A3_, potentially acting as a base to accept a proton from the bound tyrosine substrate (18). The increased flexibility of copper coordinating His residues (H_A2_, H_B1_ and H_B2_) and their role as deprotonating bases was further corroborated by measuring the increase in monophenolase activity of various of *Cg*AUS mutants (19). Our group recently discovered a novel fungal PPO (*Tt*PPO) from the ascomycete fungus *Thermothelomyces thermophila,* which is homologous to *Ao*CO4 (45% sequence identity for 82% coverage). Even though *Tt*PPO could not oxidize tyrosine, it presented high activity on other monophenolic substrates like guaiacol, hydroquinone and 1,8-dihydroxynaphthalene. Additionally, it was able to oxidize and remove various chlorophenols. Site-directed mutagenesis was performed on the proposed key activity determinants and the effect on chlorophenols removal was studied (20). In the present work, we evaluate the effect of these mutations on the kinetic characteristics of *Tt*PPO variants and discuss it based on the crystal structure of one of these variants (*Tt*PPO-GN), determined at 1.55 Å resolution. The crystal structure of *Tt*PPO is the second structure of an ascomycete PPO belonging to the short tyrosinase-family, and thus provides further evidence regarding the structure-function relations of these promiscuous enzymes.

## 2. Results and Discussion

PPOs are enzymes of high biotechnological interest, since they are used in the fields of biocatalysis, biosensors and bioremediation, while their inhibition has been widely studied, due to their implication in post-harvesting browning of fruits and mushrooms, as well as in human melanogenesis (21–27). Even though several PPOs from various sources (mostly plants) have been biochemically and structurally characterized, the molecular determinants of the monophenolase/diphenolase activity ratio have been a long-lasting debate the past decade. In the present work, we attempted to get a deeper understanding of these enzymes’ catalytic behavior, by combining mutational, biochemical and structural studies. To this end, we focused on a novel PPO (*Tt*PPO), that had been previously reported as potential bioremediation agent (20). Selection of mutation points was performed based on proposed activity determinants as analyzed in the “Introduction” section. These point mutations have been shown to increase the monophenolase activity of PPOs in some cases (14, 28, 29). The corresponding residues in *Tt*PPO are Leu306 (gate residue), Gly292 (residue H_B1_+1) and Tyr296 (residue H_B2_+1) (Fig. 1), and the produced variants involved the following mutations: G292N, L306A, Y296V, G292N/L306A and G292N/Y296V.

**Figure 1.**
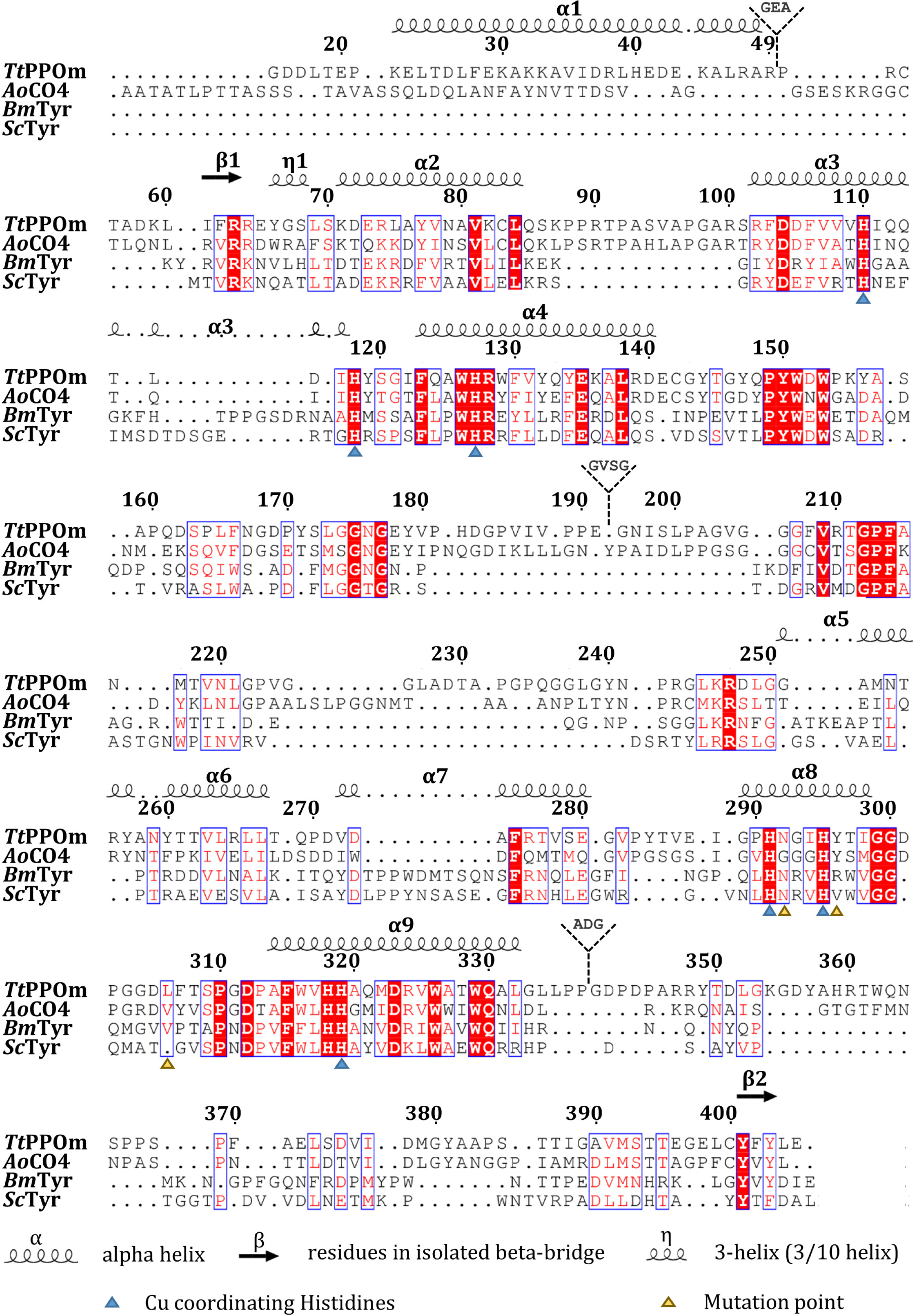
Structure-based sequence alignment of *Tt*PPO-GN, *Ao*CO4 (PDB ID: 4J3R), *Bm*Tyr (PDB ID: 3NM8) and *Sc*Tyr (PDB ID: 1WX2). Copper coordinating histidines are indicated by a blue triangle (CuA). The residues that have been mutated are denoted by a yellow triangle. The residues that were not included in the structural model are denoted as insertions. Identical and similar residues are printed in white on a red background and in red on a white background, respectively. The figure was prepared using ESPript (ESPript - http://espript.ibcp.fr) (53).

### 2.1 Substrate specificity of TtPPO variants

As previously reported (20), *Tt*PPO shows activity on some monophenolic substrates like guaiacol, hydroquinone, vanillin, 1,8-dihydroxynaphthalene, cresols and tyrosol. However, since it cannot oxidize L-tyrosine it cannot be categorized to the family of tyrosinases, based on the standing nomenclature. Hence, we could say that *Tt*PPO is a CO with limited monophenolase activity, especially on small monophenols.

Monophenolase/diphenolase ratio for all mutants, measured either on phenol/catechol or 4-chlorophenol/4-chlorocatechol, was reduced or unaltered compared to WT (data not shown). Even though these mutations did not succeed in substantially increasing the enzyme’s activity on monophenolic substrates, they appear to affect the substrate specificity of the enzyme. Kinetic analysis of the most efficiently oxidized substrates by *Tt*PPO and its variants clearly shows that the mutated residues play an important role in binding of the substrates (Table 1). Considering catechin, which is a bulky polyphenol, all variants showed improved affinity compared to the WT (3-12 times decreased *K*_M_); however, the turnover number (*k*_cat_) decreased 3-558 times as well. More specifically, mutant G292N was almost inactivated, due to an extreme decrease of *k*_cat_ combined with a moderate increase of affinity (7.8 times). On the contrary, L306A showed 3-fold increased efficiency (*k*_cat_/*K*_M_) due to considerable increase of affinity (12 times) and a more moderate decrease in *k*_cat_ (4 times). Mutant Y296V presented similar efficiency with the WT due to a proportional decrease of both *k*_cat_ and *K*_M_. Concerning double variants, both bearing the devastating G292N mutation, G292N/L306A only reduced its efficiency 2-fold compared to single variant L306A, while G292N/Y296V seemed to be more affected by the G292 mutation, reducing its efficiency by 18-fold compared to single (Y296V) variant. Based on the above, it can be seen that the size of the gate-keeper residue is critical for the accommodation of bulky substrates.

**Table 1.** Kinetic constants of wild-type (WT) *Tt*PPO and its mutants on phenolic substrates. *k_cat_* is expressed in min^-1^, *K_M_* in mM and *k*_cat_/*K*_M_ in min^-1^ mM^-1^. Values in parentheses are the standard deviation of 3 measurements.

| Substrate |  | WT | G292N | L306A | Y296V | G292N/<br>L306A | G292N/<br>Y296V |
| --- | --- | --- | --- | --- | --- | --- | --- |
| Catechol | $k_{cat}$ | 8.6(0.6) | 5.4(0.2) | 26.2(3.0) | 4.1(0.1) | 15.4 (0.5) | 4.4(0.6) |
| | $K_M$ | 32.5(4.3) | 8.1(0.9) | 7.5(1.3) | 33.9(2.3) | 5.0(0.4) | 12.56(3.0) |
| | $k_{cat}/K_M$ | 0.3(0.0) | 0.7(0.1) | 3.5(0.7) | 0.1(0.0) | 3.1(0.3) | 0.35(0.1) |
| 4-Chlorocatechol | $k_{cat}$ | 324.8(8.6) | 17.6(0.4) | 33.9(0.9) | 32.2(0.9) | 26.2(0.3) | 82.2(3.7) |
| | $K_M$ | 1.8(0.1) | 3.5(0.2) | 0.4(0.0) | 2.2(0.2) | 0.5(0.0) | 4.3(0.3) |
| | $k_{cat}/K_M$ | 180.4(11.9) | 5.0(0.3) | 81.6(6.0) | 14.6(1.2) | 48.3(2.0) | 18.9(1.7) |
| L-DOPA | $k_{cat}$ | 4.3(0.5) | 7.5(1.0) | 4.0(0.2) | 2.1(0.1) | 235.1(81.1) | 1.0(0.1) |
| | $K_M$ | 2.4(0.7) | 6.2(1.4) | 4.6(0.5) | 1.0(0.1) | 37.7(14.7) | 0.6(0.1) |
| | $k_{cat}/K_M$ | 1.8(0.5) | 1.2(0.3) | 0.9(0.1) | 2.1(0.3) | 6.2(3.3) | 1.6(0.4) |
| Catechin | $k_{cat}$ | 1619.1(549.2) | 2.9(0.3) | 415.0(35.1) | 547.3(84.6) | 337.9(11.2) | 33.3(1.7) |
| | $K_M$ | 33.7(14.0) | 4.3(1.0) | 2.8(0.5) | 9.7(2.4) | 4.3(0.3) | 10.3(0.8) |
| | $k_{cat}/K_M$ | 48.1(25.8) | 0.7(0.2) | 147.2(30.6) | 56.2(16.6) | 78.9(6.1) | 3.2(0.3) |
| Guaiacol | $k_{cat}$ | 39.3(2.2) | 6.6(0.3) | - | 12.8(0.9) | 0.5(0.0) | 1.5(0.1) |
| | $K_M$ | 4.6(0.6) | 12.4(1.5) | - | 13.3(1.9) | 6.9(1.7) | 27.3(4.5) |
| | $k_{cat}/K_M$ | 8.6(1.3) | 0.5(0.1) | - | 1.0(0.2) | 0.07(0.0) | 0.06(0.0) |
| Hydroquinone | $k_{cat}$ | 67.5(3.9) | 61.3(2.9) | 66.5(1.9) | 12.3(0.5) | 40.3(3.9) | 34.1(3.4) |
| | $K_M$ | 59.1(6.0) | 49.3(4.3) | 21.6(1.6) | 22.8(2.5) | 75.3(4.7) | 103.0(14.4) |
| | $k_{cat}/K_M$ | 1.1(0.1) | 1.2(0.1) | 3.1(0.2) | 0.5(0.1) | 0.5(0.1) | 0.3(0.1) |

L-DOPA was the only catecholamine tested, bearing a long substitution at the *p*- position of the ring. All variants showed similar catalytic efficiency compared to the WT, except for variant L306A which had a lower *k*_cat_/*K*_M_, due to reduced affinity for the substrate. Double mutant G292N/L306A, on the other hand, showed over 3 times increased catalytic efficiency. Even though its affinity for the substrate decreased 15-fold, this variant showed a remarkable increase in its turnover number, that was not witnessed for the two single variants, suggesting that the combinational mutation of these two positions is rather beneficial for the efficiency on L-DOPA.

For the WT as well as for all the variants, efficiency on 4-chlorocatechol was higher (7-600 times) than on unsubstituted catechol. However, all mutants showed decreased catalytic efficiency on 4-chlorocatechol compared to the WT (2-36 times), mostly due to significant reduction of *k*_cat_ (4-18 times). Concerning catechol, on the other hand, single mutants G292N and L306A showed an increase in catalytic efficiency by 2- and 12-fold, respectively, while Y296V’s efficiency was decreased 3 times. Double variants showed *k*_cat_/*K*_M_ values in-between the ones of each single variant. Similarly to the previous observations on catechin, the kinetic constants of double variant G292N/L306A were very similar to L306A, indicating that an alteration in the nature of the gate residue affects more significantly the catalytic behavior of *Tt*PPO.

The two tested phenolic derivatives were guaiacol, which is an *o*-methoxyphenol and hydroquinone, which is a *p*-hydroxyphenol. All mutations appeared to be devastating for the activity on guaiacol, with single mutants G292N and Y296V retaining almost the same *k*_cat_, but with a concurrent decrease in affinity. Especially for mutation L306A, kinetic constants could not be calculated, since the activity was barely detectable, implying that this residue may play an important role in binding of the substrate. On the contrary, this mutation proved to be rather beneficial for the catalytic efficiency (3-fold increase) on hydroquinone. Mutant G292N showed very similar kinetic characteristics with the WT. While the other two single variants showed decreased *K*_M_ (almost 3-fold), double variants’ affinity for this substrate was decreased, with a concurrent decrease in *k*_cat_. Mutation Y296V seems to have the most pronounced -negative- effect on *Tt*PPO activity and affinity fot hydroquinone.

Specific activity measurements with vanillin, which is a *p*-substituted guaiacol showed that all variants lose most of their activity as with guaiacol. Variants containing mutation L306A showed superior activity on epinephrine, pyrogallol and caffeic acid, while variant G292N only on the latter substrate. Interestingly, even though the WT enzyme and variant G292N did not show any activity on gallic acid, the rest of the variants did (Fig 2). Other than that, all variants were also tested in overnight reactions for phenolic substrates to which they showed lower activity, with the exception of L-tyrosine and *p*-hydroxyphenylacetic acid. Contrary to WT, all variants were active on these compounds, especially single variants L306A and Y296V. Interestingly, the WT enzyme had some activity on D-tyrosine, which increased 5-fold with mutation G292N. Most variants showed decreased activity for the small phenols that the WT enzyme could oxidize, such as *o*- and *p*-cresol, *p*-hydroxybenzoic acid and tyrosol. Exception was L306A mutant, which showed enhanced activity on tyrosol (almost 4 times) and *p*-cresol (almost 2 times), and mutant Y296V on *p*-cresol (1.5 time). Other results worth mentioning is the enhanced activity of all variants against resorcinol, with the most pronounced (5-fold) increase, being observed in variant G292N (Fig S1).

**Figure 2.**
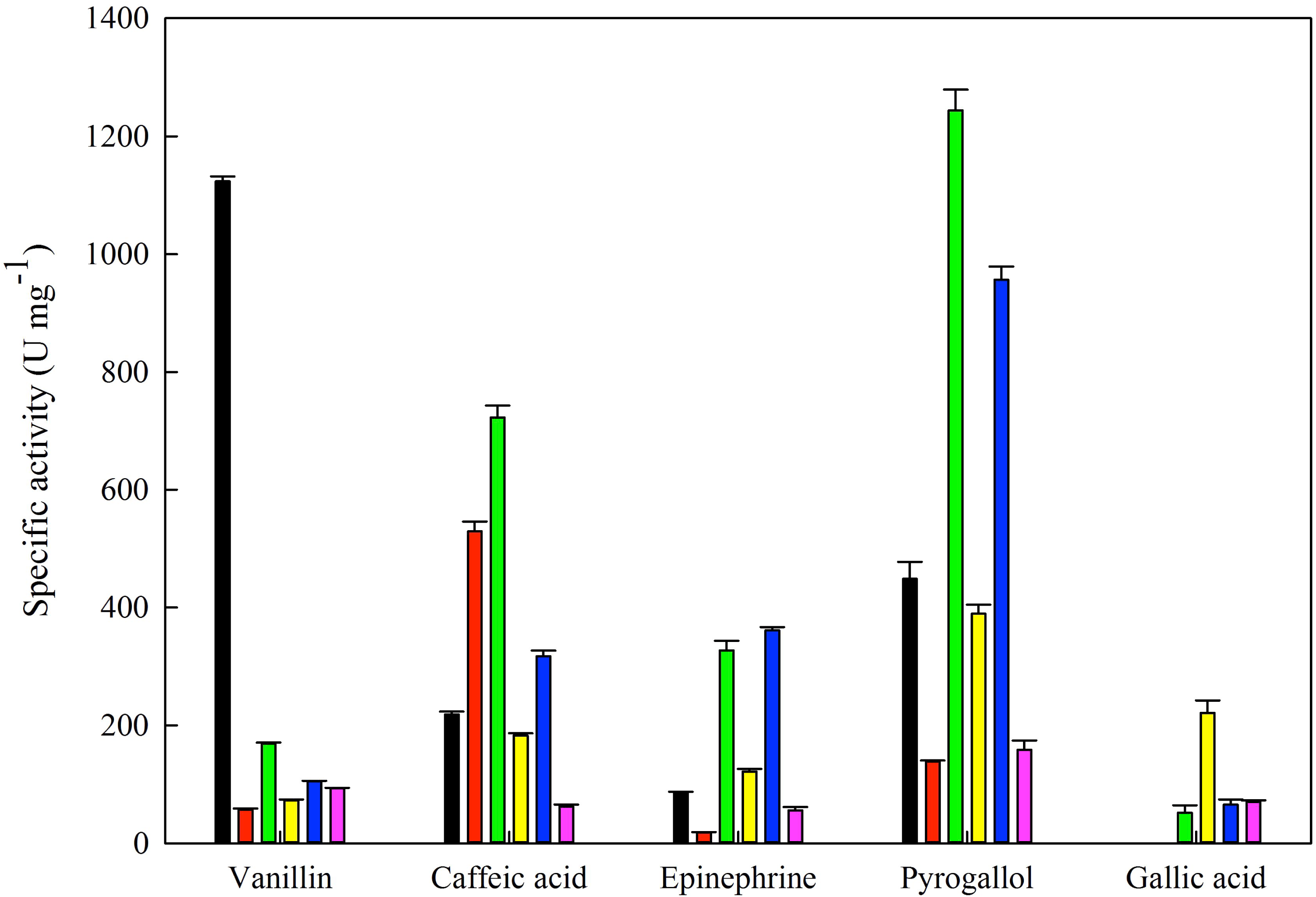
Specific activity of *Tt*PPO (black) and its variants G292N (red), L306A (green), Y296V (yellow), G292N/L306A (blue) and G292N/Y296V (magenta) measured at 40 °C. Error bars represent the standard deviation of triplicates.

What can actually be deduced from the substrate specificity experiments in the present study is that no specific pattern was detected for each of the variants, even though the targeted amino acids clearly influence enzyme specificity. Variants containing mutation L306A presented enhanced activity on many substrates including monophenols, in most cases due to increased affinity (lower *K*_M_, Table 1). This could be due to the smaller Ala side chain and thus a more “spacious” active site, however, crystallographic studies of *Tt*PPO WT and variants in complex various substrates are needed to corroborate this assumption. The implication of this residue in substrate binding has been shown in the case of dandelion PPOs, where it was proposed that a Phe at this position contributes to substrate binding via by π stacking interactions, also involving H_Β2_ (30). In order to get a better understanding of the possible enzyme-substrate interactions, the determination of *Tt*PPO crystal structure was pursued.

### 2.2 Crystal structure of TtPPO-GN (variant G292N)

*Tt*PPO-GN structure was determined 1.55 Å resolution. According to *DSSP* calculations (32, 33), *Tt*PPO-GN structure involves 9 *α*-helices, in which ∼34% of the observed amino acids (130 out of 381) participate. These *α*-helices are equally distributed in two amino acid sections: α1 to α4 are located between amino acids 22 and 140, while, α5 to α9 are located between amino acids 251 and 332 (Fig. 1). As already observed for the *Ao*CO4 structure, a four-helix bundle (α3, α4, α8 and α9) containing the copper-coordinating His, form the catalytic site of the protein (Fig. 3A). Interestingly, just after α3 and α8 and right before α4 and α9 helices, only a small loop of few amino acids (5 and 15, respectively) interferes, dividing the former helices into two partially linked helices (α3-α4 and α8-α9). The area between helices α1 and α2 (amino acids 49-70) mainly consists of a bulged loop separated in three sections by a strand (a.a 61-63) and a 3/10 helix (a.a 66-68) (Fig. 1).

**Figure 3.**
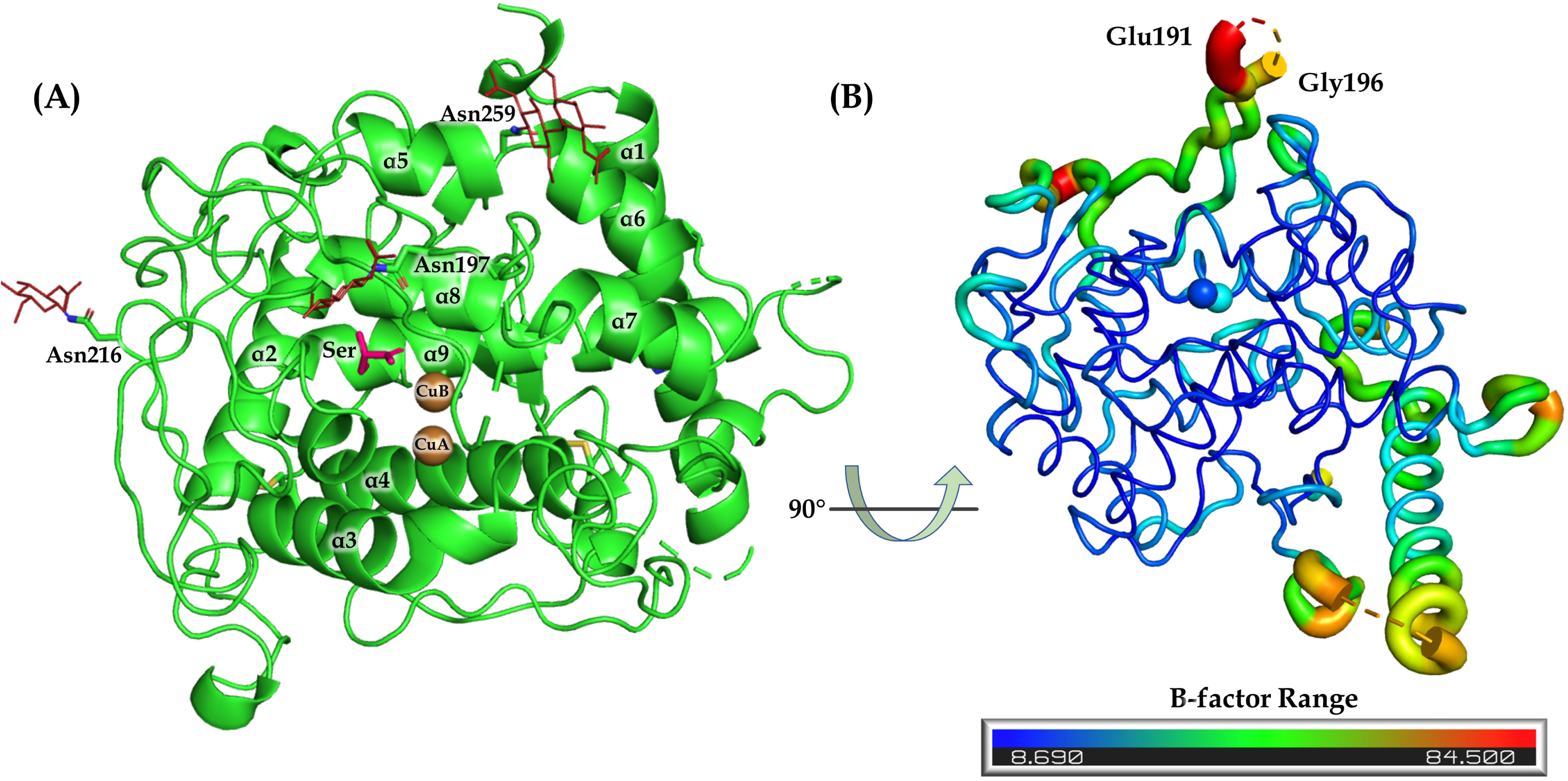
(A) Ribbon representation of overall *Tt*PPO structural model. (B) B-factor analysis of *Tt*PPO-GN.

Fourteen N-terminal (Glu1 to Val14) as well as fourteen C-terminal (Gln406 to Asp419) residues were not visible in the electron density map, while Gly50, Glu51, Ala52, Gly192, Val193, Ser194, Gly195, Ala338, Asp339 and Gly340, were also excluded from the structure due to insufficient density. Based on MW calculations, the crystallized species should be full length *Tt*PPO-GN, corresponding to the higher MW band observed in SDS-PAGE electrophoresis (Fig. S2A). Missing residues 192-195 belong to an elongated loop, located between helices a4 and a5 and shielding the copper containing active site (Fig. 1 and 3B). The relevant loop in homologous enzymes has been previously shown to allow substrate access to catalytic copper ions via its rearrangement (5). The mobility of this loop is also observed in *Tt*PPO-GN structure, as evidenced by the relatively higher B-factors of its residues (Fig. 3B), further confirming its suggested implication in the entrance of substrates to the active site.

In accordance with the predicted glycosylation sites performed in previous study (20), the crystal structure of full-length *Tt*PPO revealed three *N*-glycosylation sites: Asn197 (one *N*-acetylglucosamine-NAG), Asn216 (NAG) and Asn259 (NAG-NAG) (Fig. 3A). On the basis of the electron density, an additional residue (*β*-D-mannose-BMA) is probably present on Asn259 glycosylation chain, however, map quality in this specific area did not allow for successful modeling of this additional sugar. The overall structure is stabilized by two disulfide bonds: Cys55–Cys400, which connects and stabilizes the N- and C-termini of the polypeptide chain and Cys83–Cys142, which connects the α2 helix with an area just outside α4 helix in the active site.

### 2.3 Dicopper site analysis

The two copper ions in the catalytic center of *Tt*PPO-GN are coordinated by three histidine residues each: His110/H_A1_ (α3 helix), His118/H_A2_ (loop before α4) and His127/H_A3_ (α4) coordinate CuA while His291/H_B1_ (α8), His295/H_B2_ (α8) and His319/H_B3_ (α9) coordinate CuB, with His118 being the only residue not participating in an *α*-helix motif. Additional density between the two copper ions was modeled as a bridging water molecule (H_2_O870). The distance of this water molecule is 1.9 Å from both CuA and CuB, while the distance between the two copper ions is 3.71 Å (Fig. 4Α).

**Figure 4.**
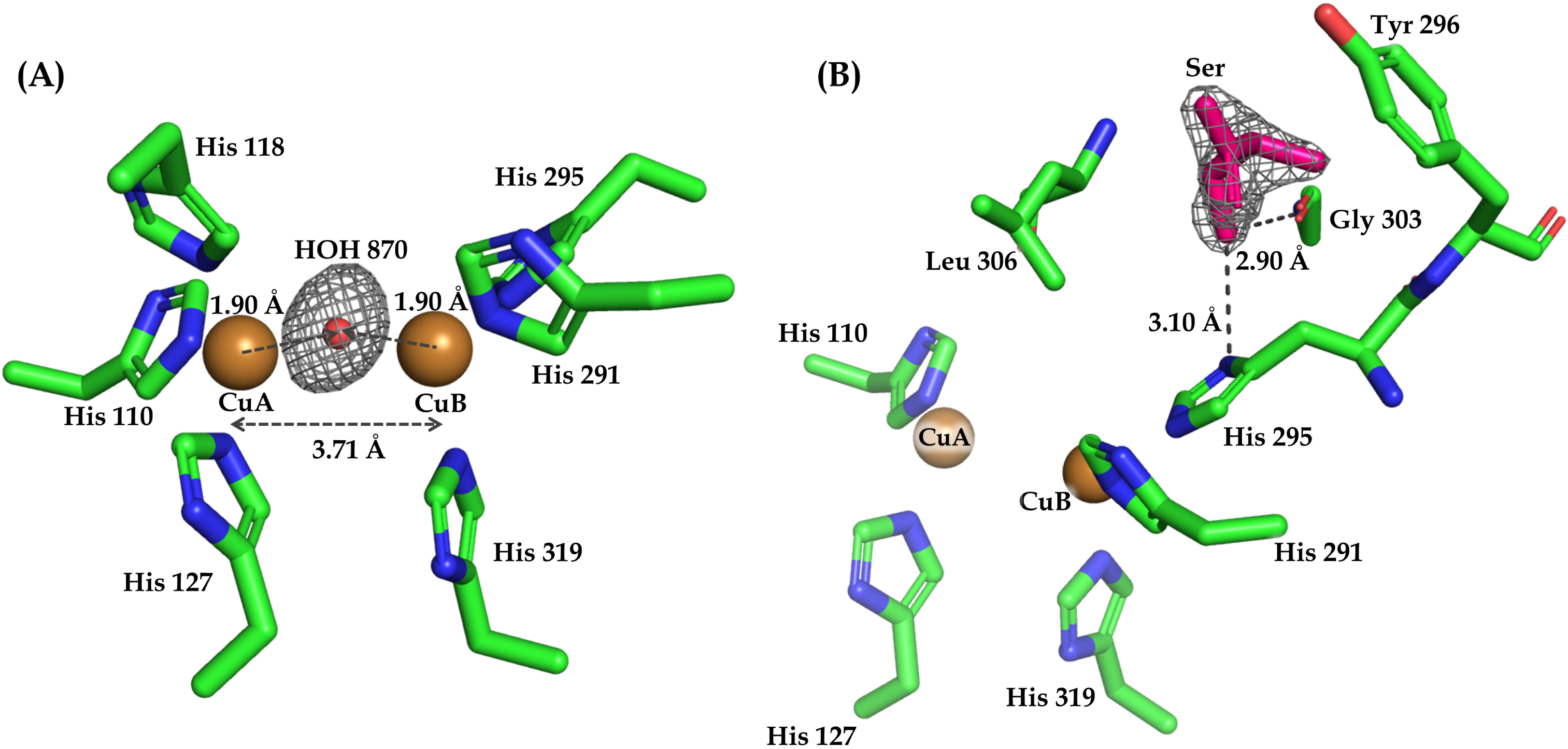
(A) Copper site in the crystal structure of *Tt*PPO-GN. An *F_o_* – *F_c_* omit map for the water molecule between the copper ions is shown as grey mesh at 5σ contour level. (B) View of the serine molecule, located on top of the dicopper center with its *2F_o_* – *F_c_* electron density map (grey mesh) contoured at 1σ. Ηydrogen bonds are denoted as grey dashed lines and the corresponding atomic distances are marked in Å.

There is an ongoing discussion concerning the exact nature of this additional electron density, evident at several structures of coupled binuclear copper (CBC) enzymes, which is directly related with the oxidation state of their active site. In the *oxy*-form, the two Cu^2+^ ions are bridged by a dioxygen molecule with a Cu-Cu distance of approximately 3.6 Å, while, in the *deoxy*-form, both copper ions are reduced to Cu^+^ state with their distance increasing to 4.6 Å. Under aerobic conditions, the *deoxy*-form quickly falls to *oxy*-form by dioxygen binding. Concerning *met*-form, it has been suggested that it consists of two Cu^2+^ ions at a distance of 3.4 Å bridged by one endogenous ligand (such as hydroxide) (7). In *Tt*PPO-GN, based on the distance between the two copper ions and the observed electron density, one could speculate that the active site of the enzyme is in *met*-form. However, lack of spectroscopic data, as well as the overall resolution of the crystallographic structure, hinder identification of the existing state.

Before data collection, *Tt*PPO-GN crystals were soaked in mother liquor supplemented with various substrates, however, no relevant electron density was observed in their active site. Instead, all crystals had an additional density on top of the dicopper center and close to Tyr296, which was modeled as serine; an ingredient of “amino acids mix” of crystallization solution. This additional amino acid seems to be firmly stabilized in this area *via* hydrogen bonds with H_B2_ (3.10 Å), and the carbonyl oxygen of Gly303 (2.90 Å) (Fig. 4Β). Tyr296 has been underlined to be critical for successful substrate orientation in the active site (11), thus serine’s location and tight binding may be the reason of the unsuccessful detection of tested substrates to the active site. A close inspection of *Tt*PPO-GN active site, with the bound serine molecule, also reveals the stereochemical constraints induced by the “bulky” Leu 306, positioned on top of CuA, when it comes to substrate access to the catalytic dicopper center (Fig. 4Β). At this point, perhaps it is worth mentioning the recent work by Biundo et al (35), who proved that some plant and mushroom PPOs could present proteolytic activity, cleaving a particular esterase’s extended loop between residues Ser and Ile, which is not a typical protease cleavage site. Even though the mechanism of this PPO activity is unclear, it points out the affinity of their active site with serine residues.

### 2.4 Comparison with other PPO structures

A search for structural homologues using the Dali server (36) revealed that *Tt*PPO-GN is most similar to *Ao*CO4 (PDB code 5OR3), with an rmsd of 1.6 Å, a Z-score of 51.9 and 43% sequence identity. This similarity is also reflected in the substrate specificity exhibited by the two PPOs, since both are active on catechol, caffeic acid, catechin, phenol, tyrosol, *p*-cresol and guaiacol (20, 37). The second and third closest structural homologues are of almost equal similarity to *Tt*PPO-GN: *Bacillus megaterium* tyrosinase (*Bm*Tyr, PDB code: 6QXD, 31% sequence identity, an rmsd of 2.1 Å and a Z-score of 27.7), followed by *Streptomyces castaneoglobisporus* tyrosinase (*Sc*Tyr, PDB code: 2ZMZ, 31% sequence identity, an rmsd of 2.1 Å and a Z-score of 26.4).

The overall structure of *Tt*PPO-GN is very close to that of *Ao*CO4, with the main differences observed in loop regions located quite distant from the active site (Fig. 5A). *Tt*PPO-GN displays a longer conformation in the loop region just after helix a9 (residues 335-346, Fig. 1). Another difference is observed in part of the flexible loop in the vicinity of the copper coordinating site, comprising residues 224 to 236 in *Tt*PPO-GN. The relevant region in *Ao*CO4 has an insertion of several amino acids (residues 216-224) and is more tilted towards the active site (Fig. 5A, highlighted in a blue frame). Τhe two small *β*-strands in *Ao*CO4 that are located between helices α4 and α5, are not observed in *Tt*PPO-GN structure. Furthermore, there are three disulfide bridges in *Ao*CO4 structure, two of which are also present in *Tt*PPO-GN. Even though the third disulfide bond which is absent in *Tt*PPO-GN has been suggested to stabilize two long loops in *Ao*CO4, the two enzymes share similar thermostability, as shown by previously published biochemical data (20).

**Figure 5.**
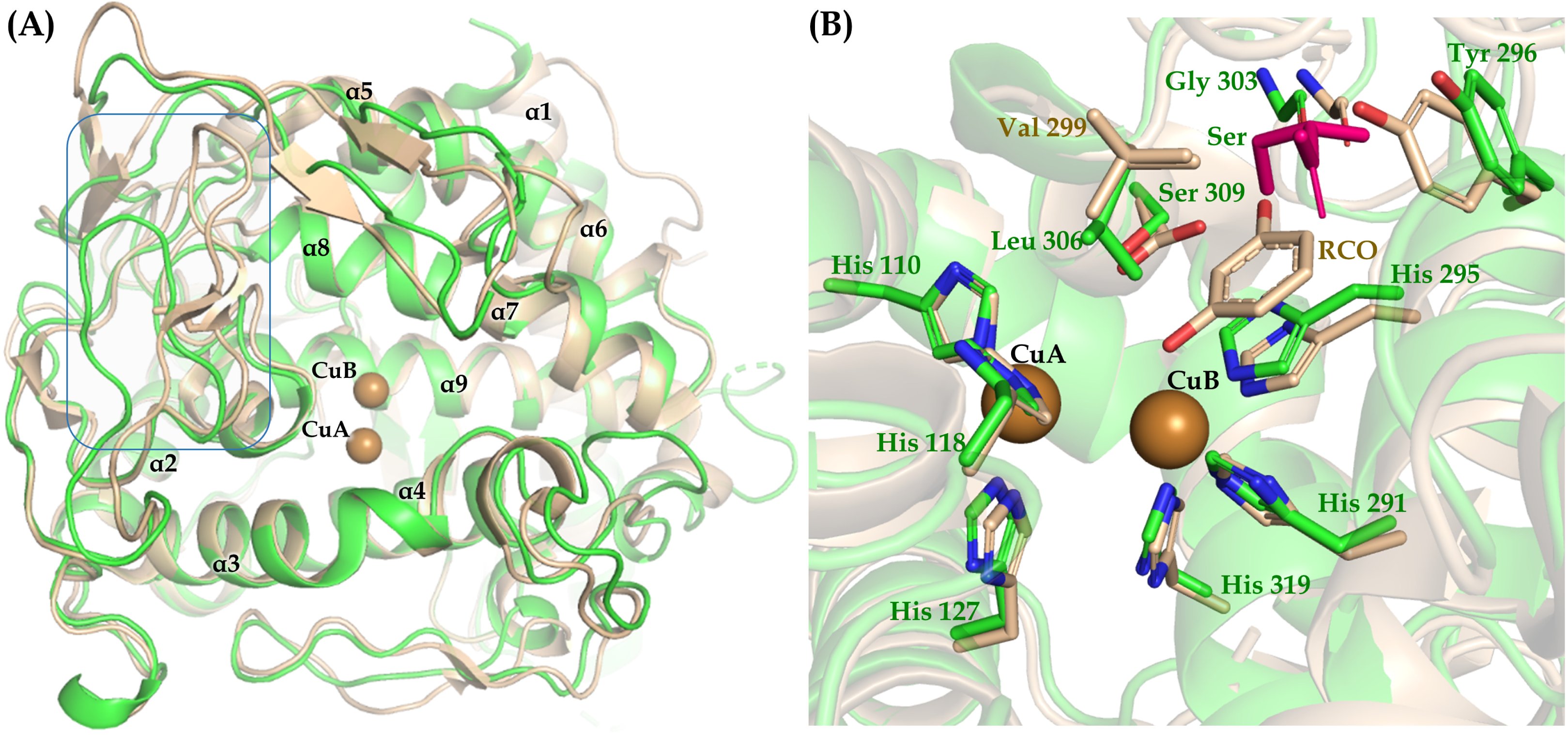
(A) *Tt*PPO-GN structure (in *green*) superimposed on to *Ao*CO4 (PDB code 6GSG, *beige*). The loop region comprising residues 224 to 236 of *Tt*PPO is highlighted in a blue frame. (B) Superposition of the copper sites of the *Tt*PPO-GN and *Ao*CO4 (with resorcinol bound, PDB code 6GSG). The residues implicated in copper and substrate coordination are shown in stick representation.

Superimposition of *Tt*PPO-GN structure with *Ao*CO4 in complex with resorcinol (PDB code 6GSG) showed that the bound serine in the former is positioned slightly above the ligand observed in the active site of its closest structural homologue (Fig. 5B). Resorcinol in *Ao*CO4 structure is coordinated by the side chain of Ser302 and the carbonyl oxygen of Gly296. Both residues are conserved in *Tt*PPO-GN (Ser309 and Gly303), suggesting a possible implication in the binding of the substrate in its active site. In addition to that, the superimposed structures clearly show that the bulkier “gate” residue in *Tt*PPO-GN (Leu instead of Val in *Ao*CO4), results in a more congested active site, with potential implications in substrate specificity. This is in accordance with the biochemical data, where mutation of Leu306 to alanine resulted in increased affinity (lower *K*_M_) for the majority of the examined substrates.

As mentioned before, monophenolase activity in PPOs has been tightly related with the occurrence of an Asn residue located just after H_B1_ (His291 in *Tt*PPO). Together with a conserved Glu residue positioned several residues before H_B1_, these two residues coordinate a water molecule, which is suggested to assist in deprotonation of monophenolic substrates, thus launching a tyrosinase catalytic cycle (11). The Glu residue is conserved in native *Tt*PPO, however, there is a Gly after H_B1._ This led us to estimate the effects of G292N mutation both on the structure and function of the enzyme. By comparing our structure with *Bm*TYR (PDB code 3NM8), it can be seen that the Glu residue (Glu280 in *Tt*PPO-GN) points towards a different direction, and its side chain does not contribute to the coordination of a water molecule in common with Asn292 (Fig. 6). H_2_O786, however, is coordinated by the side chain amide group of Asn292 and the main chain carbonyl of Glu280, but is unlikely to play the role of deprotonating water, due to its distant location from the suggested substrate binding site. On the other hand, the side chain carbonyl of Asn292 is at 3.7 Å distance from nitrogen ND1 of H_B1_ (His291) (Fig. 6A) and could thus form a hydrogen bond with the latter through a minimal rearrangement upon substrate binding, rendering H_B1_ a potential base for the deprotonation of incoming monophenols as recently suggested (19). This is however not in accordance with the biochemical data, which did not show any increase of the monophenolase activity of this variant. On the contrary, double variant G292N/L306A doubled its activity on phenol compared to the WT enzyme, but because its activity on catechol increased 10-fold, the ratio of monophenolase to diphenolase activity decreased. Based on the above, it could be assumed that substrate deprotonation originates from the flexible His (H_A2_ and H_A3_) that coordinate CuA and that are released upon substrate binding, thus acting as bases, as recently shown for *A. oryzae* and *S. castaneoglobisporus* tyrosinases (16–18) .

**Figure 6.**
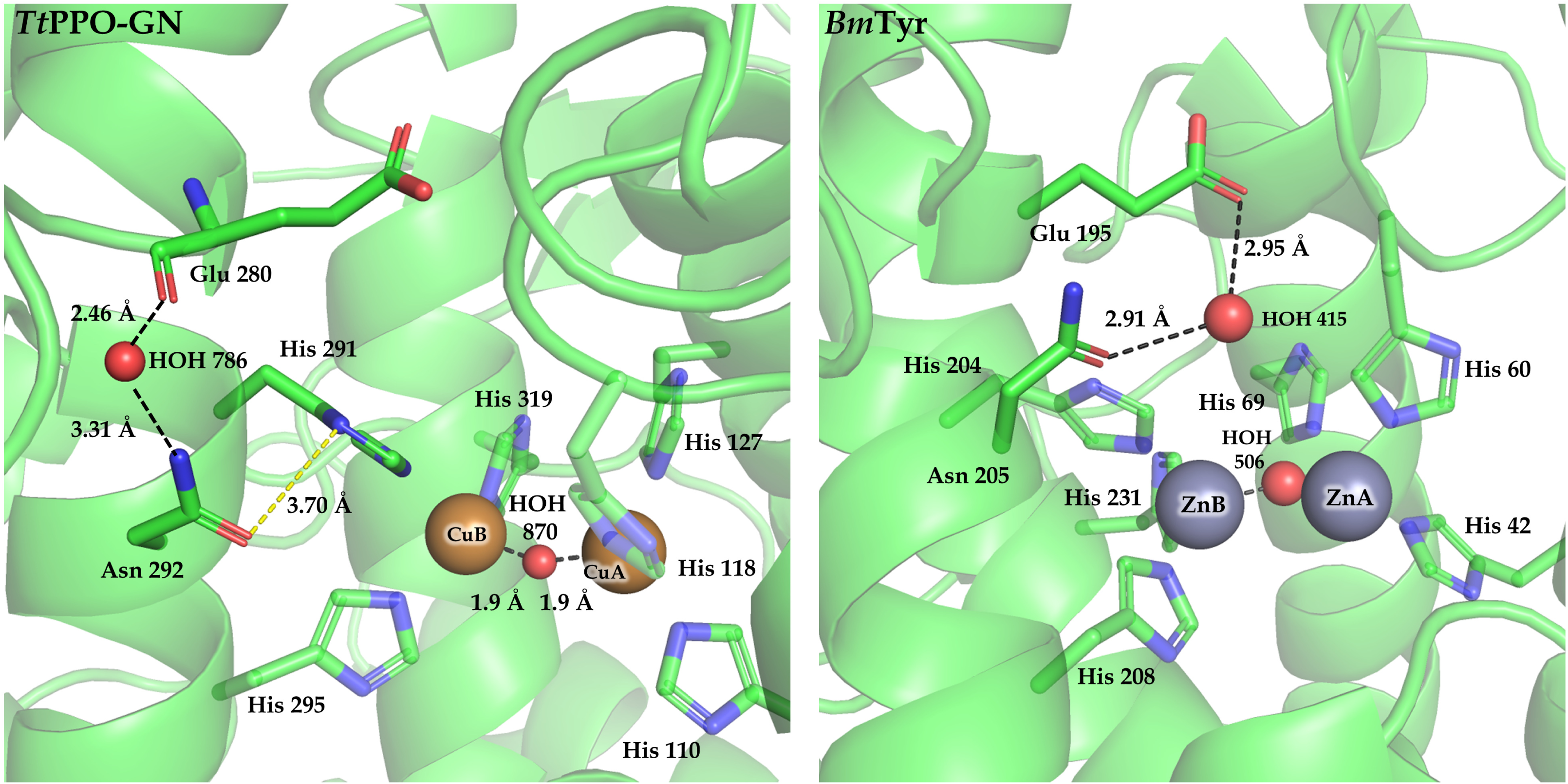
Active site of *Tt*PPO-GN (left) and *Bm*Tyr (PDB code 4P6R, right); H_B1_+1 (Asn), the conserved Glu residue and the coordinated water molecule, are highlighted. The distance between the side chains of Asn292 and H_B1_ (His291) of *Tt*PPO-GN is also shown in dashed yellow line.

## 3. Conclusions

The present work aspired to shed light on the structural determinants of PPO function, which has been a long-lasting enigma in the field, especially regarding the variability in monophenolase/diphenolase activity, exhibited by these enzymes. The biochemical characterization of *Tt*PPO mutants in combination with the crystal structure of variant *Tt*PPO-GN, resulted in the identification of amino acids in the vicinity of the copper binding site, which influence the catalytic activity of the enzyme. It is clear that the “gate” residue Leu306, positioned on top of CuA, as well as residues H_B1_+1 (Asn292) and H_B2_+1 (Tyr296) influence the catalytic activity of *Tt*PPO. The most pronounced changes were observed for the former, since its mutation to a smaller aminoacid (Leu to Ala) resulted in an increase in the activity of the enzyme against most of the tested substrates. Other than that, mutation of residue H_B1_+1 (G292N) resulted in a variant with considerably increased activity on resorcinol, D-tyrosine and *p*-hydroxyphenylacetic acid. A comparative examination *Tt*PPO-GN structure along with biochemical data, showed that residue H_B1_+1 is unlikely to contribute to substrate deprotonation as suggested from previous studies, thus pointing to a recently suggested theory where substrate deprotonation is mediated by one of the copper-coordinating His. The determination of *Tt*PPO structure in complex with its preferred substrates will allow the engineering of a biocatalyst with improved properties, given its potential use as a bioremediation agent.

## 4. Materials and Methods

### 4.1 Expression and purification of recombinant enzymes

*Pichia pastoris* recombinant strains expressing *Tt*PPO and its variants were cultivated in 500 mL buffered complex methanol medium (BMMY) supplemented with 25 μM CuSO_4_. Induction took place for 4 days at 23 ^°^C and 200 rpm by the addition of 0.5% v/v methanol per day, based on previous optimization (20). Culture broth was concentrated and dialyzed against 50 mM Tris-HCl, 300 mM NaCl pH 8 buffer and loaded on immobilized metal cobalt affinity chromatography (IMAC) column as described previously (38).

Variant G292N (*Tt*PPO-GN) that was used for crystallographic studies was expressed in 2-L cultures as mentioned above. The culture broth was concentrated and the recombinant protein was deglycosylated in the crude protein solution by the addition of 30 μg mL^-1^ EndoH (endo-*β*-*N*-acetyl glucosaminidase, EC 3.2.1.96) from *Streptomyces plicatus*. After 1.5 h incubation at 37 ^°^C the crude protein was dialyzed and purified using IMAC as above. Fractions containing the purified *Tt*PPO-GN were pooled, concentrated and applied onto a HiPrep^™^ 16/60 Sephacryl^®^ S-100 HR (GE Healthcare, Chicago, Illinois, USA) equilibrated and eluted with 20 mM Tris-HCl, 150 mM NaCl pH 8 buffer. Fractions of 2 mL were collected at a flow rate of 1 mL min^-1^ and the ones with pure *Tt*PPO-GN were pooled, dialyzed against 20 mM Tris-HCl pH 8 buffer and concentrated at 20 mg mL^-1^. The homogeneity of the protein was checked by sodium dodecyl sulfate polyacrylamide gel electrophoresis (SDS-PAGE).

### 4.2 Biochemical characterization of TtPPO variants

A typical enzymatic assay was performed, as described previously (20). Briefly, reactions containing 230 μL of 5 mM substrate in 0.1 M sodium phosphate buffer pH 7 were mixed with 20 μL of enzyme and incubated in a SpectraMax 250 microplate reader (Molecular Devices, USA) set at 40 ^ο^C. Increase in absorbance at each product’s *λ*_max_ was recorded for 20 min. One unit (U) of enzymatic activity was determined as 1 ΔΑ*_λ_*_max_ min^-1^. The substrate range of the purified enzymes and their specific activity were determined as described in (20).

Kinetic studies of the purified enzymes were performed by assaying varying concentrations of 4-chlorocatechol (0-5 mM), catechol (0-60 mM), catechin (0-10 mM), L-DOPA (0-7 mM), vanillin (0-5 mM), guaiacol (0-15 mM) and hydroquinone (0-70 mM). Kinetic constants were estimated using a non-linear regression model in GraphPad Prism 5 from GraphPad Software, Inc. (USA).

Protein amount of purified enzyme was quantified through *A*_280_ measurements(39) using a molar extinction coefficient of 75540 M^-1^ cm^-1^ calculated with ProtParam tool from ExPASy (40).

### 4.3 *Crystallization, data collection and structure determination of Tt*PPO-GN

Similarly to the native enzyme (20), SDS-PAGE analysis showed that purified *Tt*PPO-GN appears as two bands, which after deglycosylation with EndoH, appeared to be approximately 48 kDa and 39 kDa. Based on our previous analysis, the higher MW band must correspond to full length *Tt*PPO-GN, while the lower band is assumed to be an N-terminally cleaved enzyme variant. Neither IMAC nor gel filtration purification steps were successful in separating these two bands; they were thus concentrated to 10 mg mL^-1^ in 20 mM Tris/HCl pH 8.0, and submitted to crystallization experiments at the High Throughput Crystallization Laboratory (HTX Lab) of EMBL Grenoble *via* the sitting drop vapor diffusion technique (41). Crystals (Fig. S2B) appeared after 1 month in a single condition from Morpheus screen (Molecular Dimensions, condition H12: 0.1M amino acids mix, 0.1M Tris Bicine pH 8.5, 37.5% v/v precipitant mix 4) , and were prepared for X-ray data collection experiments using the CrystalDirect technology (42). Data collection was carried out at 100 K on beamline P14 of EMBL Hamburg, Germany. The crystals diffracted anisotropically, and the X-ray data were processed using the software STARANISO(43) as implemented in the autoPROC pipeline (44). *Tt*PPO crystals belong to space group *P*2_1_2_1_2, with unit-cell dimensions *a* = 86.40, *b* = 97.18 and *c* = 46.99 Å, and one molecule per asymmetric unit. The structure of *Tt*PPO-GN was solved via molecular replacement at 1.55 Å resolution using *Phaser* (45) as implemented in the *CCP4 suit* (46). The search model was derived from the crystal structure of *A. oryzae* catechol oxidase (*Ao*CO4), which shares 45% sequence identity for 82% coverage with *Tt*PPO (PDB entry 4J3R) (31). The final model was constructed by alternating cycles of refinement and model building in *REFMAC* (47) and *Coot* (48). Refinement converged to *R*_work_ and *R*_free_ values of 15.74% and 20.66%, respectively. The validation of the final refined model was performed with *PDB-REDO* (49), *MolProbity* (50) and *pdb-care* web servers (51), respectively. Atom coordinates and structure factors have been deposited in the PDB (accession code 6Z1S). Data collection and refinement statistics are summarized in Table 2.

**Table 2.** Diffraction data and refinement statistics for *Tt*PPO-GN.

| *Data collection and Refinement Statistics* |  |
| --- | --- |
| **Data collection** |  |
| Beamline | P14 (EMBL, Hamburg) |
| Wavelength (Å) | 0.98012 |
| Space group | *P* 2<sub>1</sub> 2<sub>1</sub> 2 |
| Cell dimensions (a, b, c) (Å) | a=86.40, b=97.18, c=46.99 |
| Resolution range (Å) | 43.20-1.55 (1.72-1.55)<sup>a</sup> |
| Ellipsoidal resolution (Å) (direction) | 1.44 (a*) |
|  | 1.93 (b*) |
|  | 2.24 (c*) |
| No. of molecules per asymmetric unit | 1 |
| No. of unique reflections | 30047 (1502)<sup>a</sup> |
| Completeness (%) (spherical) | 52.0 (10.1)<sup>a</sup> |
| Completeness (%) (ellipsoidal) | 86.3 (61.3)<sup>a</sup> |
| <sup>b</sup> *R*<sub>merge</sub> (%) | 15.5 (179)<sup>a</sup> |
| <sup>c</sup> *R*<sub>pim</sub> (%) | 4.4 (51.6)<sup>a</sup> |
| Mean(*I*/sd(*I*)) | 12.4 (1.8)<sup>a</sup> |
| <sup>d</sup> CC<sub>1/2</sub> | 0.999 (0.605)<sup>a</sup> |
| Multiplicity | 13.3 (12.6)<sup>a</sup> |
| Wilson *B* value (Å<sup>2</sup>) | 17.7 |
| **Refinement statistics** |  |
| *R*<sub>work</sub>/*R*<sub>free</sub> (%) | 15.74/20.66 |
| R.m.s.d., bond lengths (Å) | 0.015 |
| R.m.s.d., bond angles (°) | 1.83 |
| No. of protein atoms | 3016 |
| No. of solvent atoms | 359 |
| **Average B-values (Å<sup>2</sup>)** |  |
| All proteins | 21.70 |
| Water molecules | 30.34 |
| Copper ions | 21.23 |
| **<sup>e</sup>Ramachandran statistics (%)** |  |
| Favored | 96.78 |
| Outliers | 0 |
| Allowed | 3.22 |
| PDB ID | 6Z1S |

<sup>a</sup> Numbers in parentheses refer to the highest resolution shell, <sup>b</sup> $R_{\text{merge}} = \sum_{hkl} \sum_i |I_i(hkl) - \langle I(hkl) \rangle| / \sum_{hkl} \sum_i I_i(hkl)$ , <sup>c</sup> $R_{\text{pim}} = \sum_{hkl} (N-1)^{-1/2} \sum_i |I_i(hkl) - \langle I(hkl) \rangle| / \sum_{hkl} \sum_i I_i(hkl)$ , <sup>d</sup> CC1/2 is the correlation coefficient between two random half datasets, <sup>e</sup> Calculated using Molprobity (http://molprobity.biochem.duke.edu/)

Structure figures were prepared in *PyMol* (The PyMOL Molecular Graphics System, Version 2.3.2, Schrödinger, LLC). Structure-based sequence alignment was performed in *UCSF Chimera* (52) using the structures of *Tt*PPO-GN, *Ao*CO4 (PDB ID: 4J3R), *Bm*Tyr (PDB ID: 3NM8) and *Sc*Tyr (PDB ID: 1WX2). The alignment output was visualized using ESPript (ESPript - http://espript.ibcp.fr).

## Data availability

There are no newly determined accession numbers generated in this study. Data regarding structure determination are available at https://www.rcsb.org under the ID 6Z1S.

## Acknowledgements

This research is co-financed by Greece and the European Union (European Social Fund-ESF) through the Operational Programme «Human Resources Development, Education and Lifelong Learning 2014-2020» in the context of the project “Structural studies of a polyphenol oxidase with applications in the bioremediation of polluted waters” (MIS 5047137). The authors would like to thank the High-Throughput Crystallisation Laboratory (HTX Lab) of the EMBL Grenoble for performing the crystallization and data collection, in the frame of iNEXT project (PID 4804), which was funded by the European Community’s Seventh Framework Programme H2020 (H2020 Grant #653706). EN and MD particularly thank research technician Guillaume Hoffmann for the great cooperation.

